# A One-Step Synthesis of DNA Dumbbells Tunnels Short-Read Libraries into a Long-Read Format

**DOI:** 10.1101/2022.05.01.490243

**Authors:** Ya-Hui Chang, Hong-Hsing Liu

## Abstract

DNA dumbbells have many applications. For example, they can be templates for rolling-circle amplification to produce highly accurate long-read sequences of single molecules. We have developed a highly efficient and simple method for ‘tunneling’ short-read Illumina sequencing libraries into long-read dumbbells. A 5’-exonuclease used in Gibson Assembly converted fixed Illumina P5/P7 sequences into sticky ends. Self-looping oligonucleotides with complementary 3’-overhangs were ligated to form dumbbells. The reaction proceeded in one pot and modified only end nucleotides of input libraries. Resulting dumbbells were ready for Phi29-dependent rolling circle amplification. Demonstrative readouts of full-length 16S rDNA from a standard microbial community, obtained on a Pacific Biosciences (PacBio) platform, confirmed successful tunneling of initially P5/P7-ended sequencing libraries. Twelve fecal samples additionally showed significant correlations between standard and tunneled 16S sequence variants on a PacBio platform.

## Introduction

DNA dumbbells are a variety of circular DNA. Topologically equivalent to circles, DNA dumbbells are highly resistant to exonucleases and can serve as stable vectors for regulating gene expression^1^. Additionally, dumbbell DNA can serve as an iterative template for rolling-circle amplification^2^. Many modern sequencers use DNA dumbbells or circles as sequencing templates. Pacific Biosciences (PacBio) platforms, for example, derive long and highly accurate sequences *via* alignment of iterative ‘subreads’ into consensus ‘HiFi’ reads^3^. MGI Tech instead uses circular DNA to amplify sequencing signals *via*‘DNA nanoballs’^4^.

With high-throughput sequencing, a complete human genome can now be analyzed within hours^5^. Decreased costs and increased throughputs have accelerated application of sequencing technology to all fields of life science. However, the need to transform sequencing subjects into suitable library formats remains a key pragmatic constraint. For example, short-read libraries for Illumina are linear DNA with fixed P5/P7 sequences on either end^6^. Long-read libraries for PacBio are instead dumbbell-shaped with fixed loop sequences^7^. These libraries require different workflows to prepare. Each sequencing platform has advantages and disadvantages^8^. Illumina platforms are highly accurate but suffer from short reads of 150bp paired-end on most recent hardware. PacBio platforms instead offer long reads of well over 10kbp, achieving accuracy through iteratively optimized ‘HiFi’ consensus^3^. For some libraries, such as those that include full-length B-lymphocyte heavy chain sequences, neither platform is readily used. These libraries are typically longer than what Illumina platforms can comfortably sequence, but few established workflows exist to format these libraries for PacBio platforms. A potential solution unifies the protocols through a ‘tunneling’ option. We have realized this solution with a novel one-step synthesis of DNA dumbbells.

## Materials and Methods

### Preparation of 16S rDNA

16S rDNA of variable lengths from the DH5α *E. coli* strain were PCR amplified using the following thermocycle: 95°C 3 minutes, 30 cycles of 98°C 20 seconds + 60°C 15 seconds + 72°C 2 minutes, and 72°C 7 minutes. Each 50 μL reaction contained either a 10 μL pellet from a 5-hour culture or 10 ng template DNA, 0.2 μM each primer, and KAPA HiFi HotStart ReadyMix (Roche). A full-length founder rDNA of 1.6 kbp was prepared with the primer pair 5’-AAGCAGTGGTATCAACGCAGAGT AGRGTTYGATYMTGGCTCAG-3’ and 5’-CAGACGTGTGCTCTTCCGATCT RGYTACCTTGTTACGACTT-3’. P5 and P7-flanking derivatives of lengths varying by factors of two were subsequently amplified from the template by a common primer 5’-GGCGACCACCGAGATCTACAC AGRGTTYGATYMTGGCTCAG-3’ coupled with one of the primers 5’-GCAGAAGACGGCATACGAGAT RGYTACCTTGTTACGACTT-3’, 5’-GCAGAAGACGGCATACGAGAT CCAAGTCGACATCGTTTACGG-3’, 5’-GCAGAAGACGGCATACGAGAT CCTTCCTCCCCGCTGAAAGTA-3’, 5’-GCAGAAGACGGCATACGAGAT GCACATCCGATGGCAAGAGG-3’, and 5’-GCAGAAGACGGCATACGAGAT ACATTACTCACCCGTCCGCCACT-3’ for each template.

### Dumbbell formation

16S rDNA inputs reacted with two self-looping oligonucleotides (SLOs) in NEBuilder^®^ HiFi DNA Assembly Master Mix (New England BioLabs^®^) at 45°C for 2-3 hours. The 5’-ends of SLOs were phosphorylated and protected by five consecutive phosphorothioate bonds (*). The 3’-ends of SLOs were compatible with the corresponding end sequences of indicated inputs. Products in Fig. 1 were formed from the SLO pair Pho5’-G*G*C*C*A*GCAGC AGAGGAGGAC GCTGCTGGCC ATATG AATGATACGGCGACCACCGAGATCTACAC-3’ and Pho5’-G*A*A*G*T*GCGCTG TAAGTATTAC CAGCGCACTTC ATAT CAAGCAGAAGACGGCATACGAGAT-3’, while those in Fig. 2 were formed from the SLO pair Pho5’-G*G*C*C*A*GCAGC AGAGGAGGAC GCTGCTGGCC AAGCAGTGGTATCAACGCAGAGT-3’ and Pho5’-G*A*A*G*T*GCGCTG TAAGTATTAC CAGCGCACTTCT ATAATGG CAGACGTGTGCTCTTCCGATC-3’.

**Figure 1.**
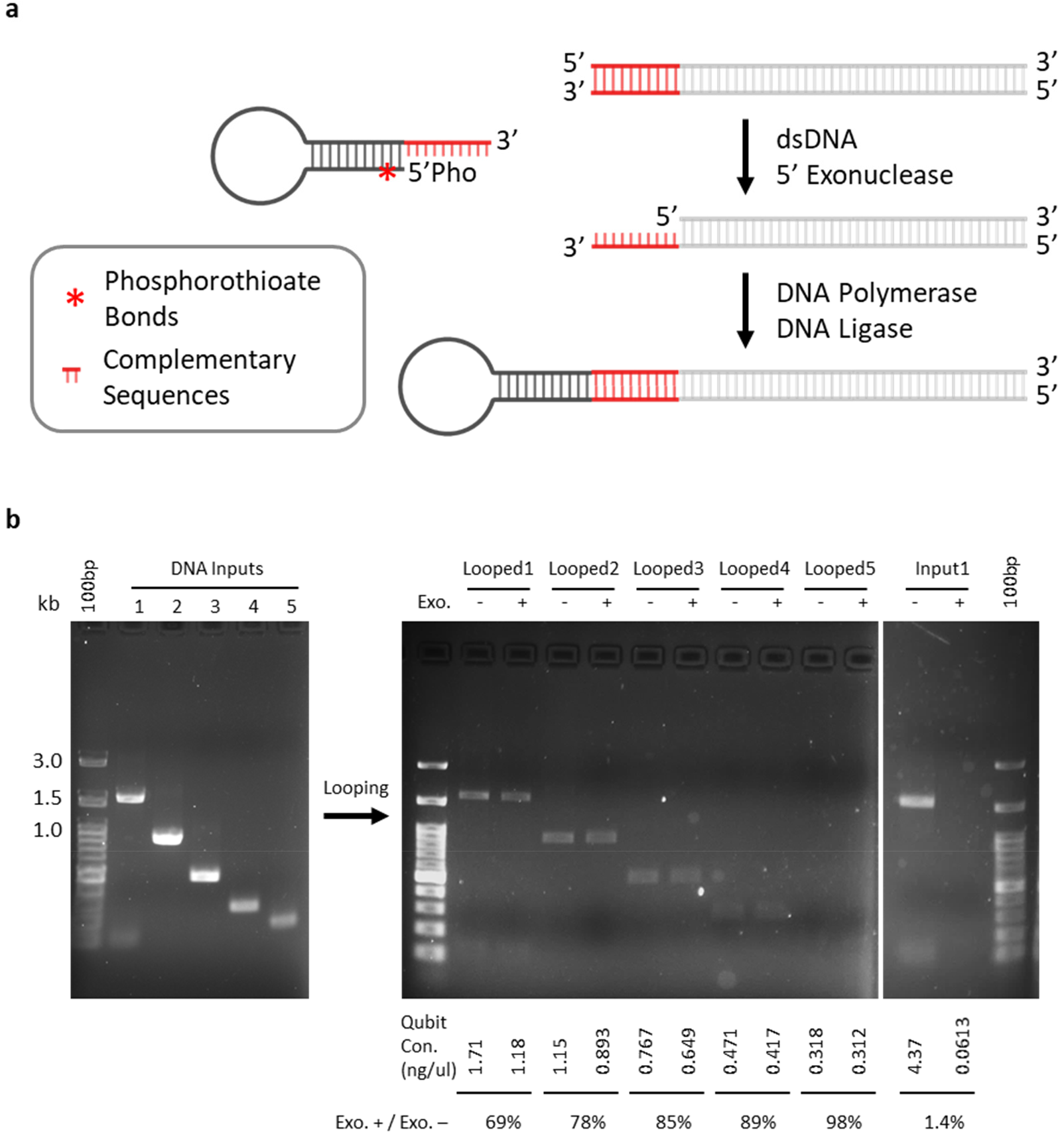
Demonstration of protocol. (**a**) P5/P7 ends of input libraries are processed using a 5’-exonuclease, as in Gibson Assembly. Only one end is illustrated. 3’-ssDNA is exposed by dsDNA 5’-exonuclease, which allows 5’-phosphorothioated and 5’-phosphorylated self-looping oligonucleotides to anneal with complementary 3’-overhangs. DNA polymerase and ligase complete the looping reaction. (**b**) Five DH5α 16S libraries of lengths varying by factors of two were subjected to conversion. Products were tested with a cocktail of exonucleases, which has no effect on successfully looped dumbbells. Conversion efficiencies varied from 69% for the longest input sequence to 98%for the shortest input sequence. Little unconverted input survived digestion by the exonuclease cocktail.

**Figure 2.**
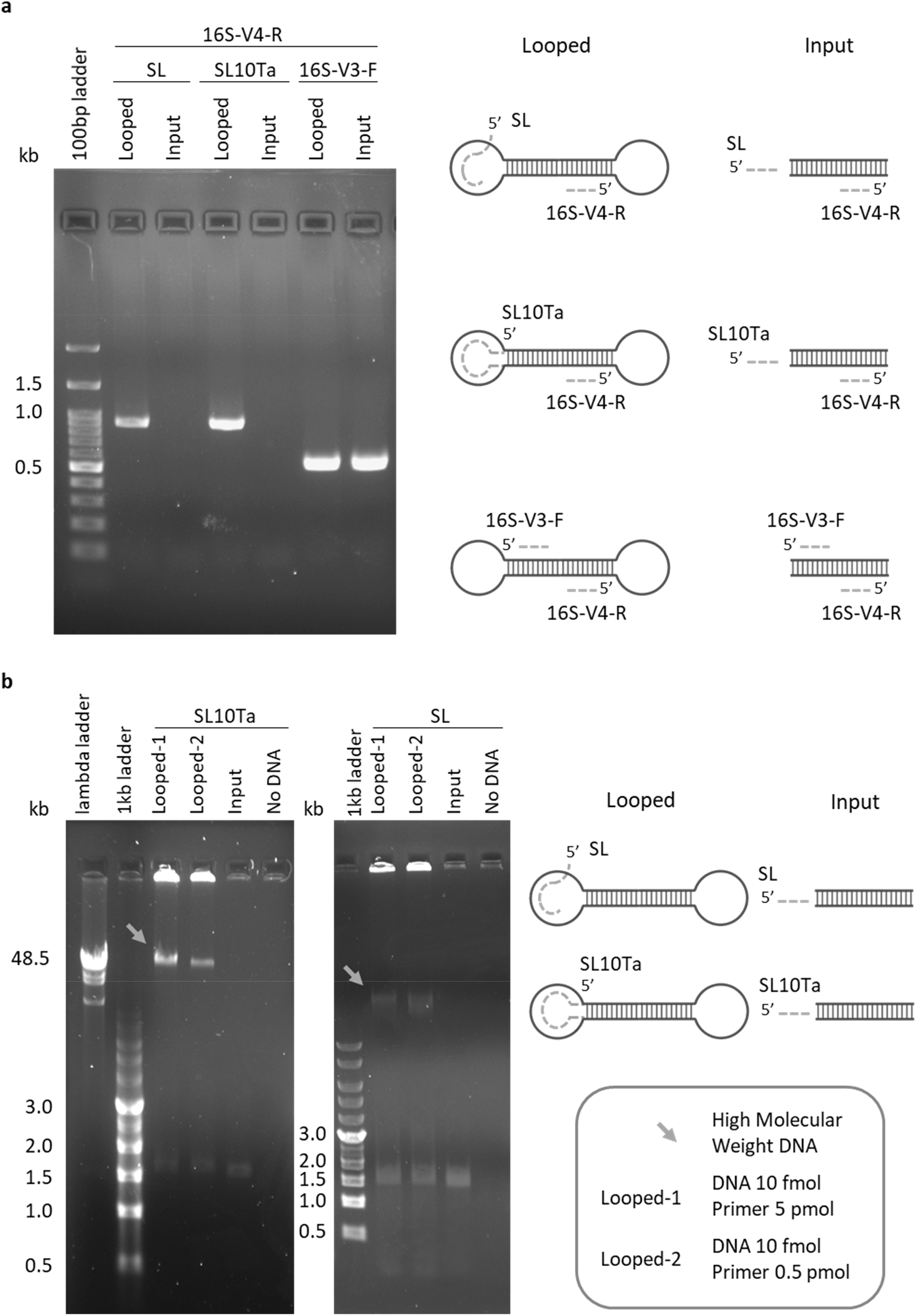
Verification of protocol. (**a**) Two different primers capable of hybridizing with a loop paired with a reverse primer matching the input sequence. Only looped products, and not unconverted inputs, yielded positive PCR bands. (**b**) Rolling circle amplifications by Phi29 polymerase proceeded only on dumbbell templates primed with two different loop-hybridizing primers. No high molecular-weight products (arrow) were produced using unconverted inputs as templates.

### Efficiency calculation

Successfully formed dumbbells resist exonuclease digestion, but DNA inputs with open ends do not. To calculate efficiency, one half of each cleanup prepared using MinElute PCR Purification Kit (QIAGEN) was treated with a cocktail of exonucleases comprising 0.2 U/μL T7 exonuclease, 0.2 U/μL truncated exonuclease VIII, 0.2 U/μL exonuclease I, and 1 U/μL exonuclease III (New England BioLabs^®^), while the other half of each cleanup was not treated. After 30 minutes at 25°C followed by 30 minutes at 37°C, remaining DNA in each half was quantified using a DNA Qubit™ dsDNA HS Assay Kit (Invitrogen™). Efficiency of dumbbell formation was derived accordingly.

### PCR verification

A common primer, 5’-GTGACTGGAGTTCAGACGTGTGCTCTTCCGATCT GACTACHVGGGTATCTAATCC-3’ or 16S-V4-R, was paired with loop-hybridizing primer SL (5’-GTCCATCCTCGTCCTCCTC-3’) or SL10Ta (5’-AGCAGC GTCCTCCTCT GCTG-3’) to verify dumbbell formation, and 16S-V3-F (5’-ACACTCTTTCCCTACACGACGCTCTTCCGATCT CCTACGGGNGGCWGCAG-3’) to control for inputs. PCR reactions were performed in Q5 HiFi polymerase ready mix (New England BioLabs^®^) using the following thermocycle: 98°C 30 seconds; 5 cycles of 98°C 10 seconds, 45°C (SL) or 62°C (SL10Ta or 16S-V3-F) 3 minutes; 25 cycles of 98°C 10 seconds, 54°C (SL) or 62°C (SL10Ta or 16S-V3-F) 30 seconds, 72°C 40 seconds; and 72°C 2 minutes.

### Rolling circle amplification

Each Phi29-dependent rolling circle amplification on dumbbells was primed with loop-hybridizing primer SL or SL10Ta. Specifically, a 10 μL mixture of 10 fmol dumbbell template, 0.5 or 5 pmol primer, and 1mM dNTP was heated at 95°C for 3 minutes before ramping down slowly to 25°C. Bovine serum albumin at final 0.2 mg/mL and Phi29 polymerase (New England BioLabs) were added and the reaction ran at 30°C for 16 hours before inactivation at 65°C for 10 minutes.

### Tunneled PacBio sequencing

Customized primers for amplifying full-length 16S with P5/P7 ends are illustrated in **Fig. 3a**. Sequence details are in **Supplementary Table 1**. PCR thermocycles were: 95°C 3 minutes, 25-28 cycles of 95°C 30 seconds + 57°C 30 seconds (ramp rate ≤ −3 °C/sec) + 72°C 1 minute, and 72°C 3 minutes. Each 25 μL reaction contained 2 ng template DNA, 0.375 μM each primer, and KAPA HiFi HotStart ReadyMix (Roche). PCR products were cleaned up with MinElute PCR Purification Kit (QIAGEN) and quantified by DNA Qubit™ dsDNA HS Assay Kit (Invitrogen™). Purified amplicons receiving the same pair of SLOs were pooled equally before assembly. Most samples were subjected to asymmetric looping (**Supplementary Table 2**), which allowed docking of the PacBio sequencing primer on the P5 loop only. One sample received symmetric looping, which took the PacBio sequencing primer on both loops. SLOs for asymmetric looping are Pho5’-C*T*C*T*C*TCAACAACAAC TCCTCCTCCTCCGTT GTTGTTGTTGAGAGAGAT GATAC GGCGACCACCGAGATCTACAC-3’ and Pho5’-G*A*A*G*T*GCGCTG TAAGTATTAC CAGCGCACTTC ATAT CAAGCAGAAGACGGCATACGAGAT-3’. The latter was replaced with Pho5’-C*T*C*T*C*TCAACAACAAC TCCTCCTCCTCCGTT GTTGTTGTTGAGAGAGAT CAA GCAGAAGACGGCATACGAGAT −3’ for symmetric looping. The assembly condition has been described above in ‘Dumbbell formation.’ Two samples were from DH5α *E. coli* strain, one sample was ZymoBIOMICS Microbial Community Standard (D6300, ZYMO RESEARCH), and others were controls that were previously sequenced on the PacBio platform.

**Figure 3.**
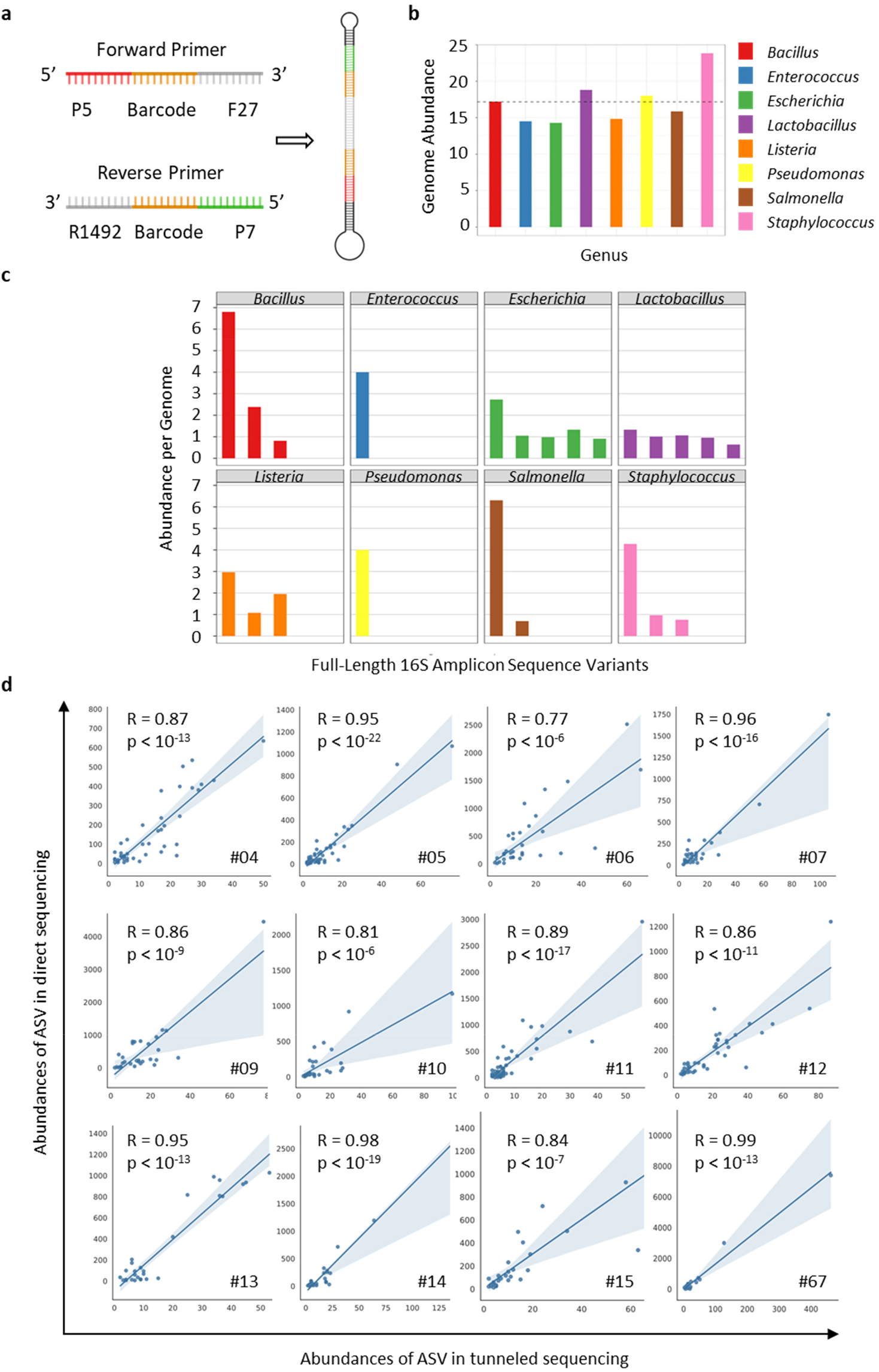
Tunneling to PacBio sequencing. (**a**) A pair of primers able to amplify full-length 16S with P5/P7 end sequences is illustrated. (**b**) Genome abundances of eight bacterial genera were relatively equal, corresponding to the known composition of the microbial community standard. (**c**) Stoichiometric analyses of 16S ASV per bacterial genome revealed integral values and sums in line with expectations. (**d**) ASV abundances are significantly correlated between direct and tunneled sequencing.

### Bioinformatic analysis of tunneled PacBio sequencing

In brief, adapters were recalled using PacBio’s utility *recalladapters.* The output subreads.bam was processed by the *ccs* utility to get the fastq file. After demultiplexing, records belonging to the standard microbial community or each fecal sample were pooled into indicated fastq files for downstream analyses. Commands and Python scripts are briefed in the supplementary file.

## Results

### One-step synthesis of DNA dumbbells

This study followed Gibson Assembly^9^ in utilizing 5’-exonucleases to expose sticky ssDNA in fixed P5/P7 sequences on Illumina libraries (**Fig. 1a**). Introduction of self-looping oligonucleotides with complementary 3’-ends completed dumbbell formation in one pot. Phosphorothioate bonds were artificially introduced at the 5’-ends of looping oligonucleotides to resist the action of 5’ exonucleases.

As a demonstration of the method, five sequencing libraries of DH5α 16S with P5/P7 ends were prepared, with lengths varying over four powers of two (**Fig. 1b**, left panel). Library conversions were performed using commercially-available ready-mix NEBBuilder kits. Purified products were subjected to digestion by a cocktail of exonucleases (**Fig. 1b**, right panel). Only successfully converted dumbbells were not degraded by the exonuclease cocktail. Conversion efficiency decreased moderately with input length, ranging from 69% for the longest input to 98% for the shortest input. Without conversions, input DNA was highly susceptible to exonucleases; only 1.4% of unconverted input DNA remained undegraded in the longest input (**Fig. 1b**, right panel).

### Verification of dumbbell formation

Looping reactions were verified using PCR (**Fig. 2a**). Dumbbells with known loop sequences were prepared from the DH5α 16S sequence. Two primers, SL and SL10Ta, differ in end sequences but hybridize with the same loop of each dumbbell. Both were each paired with a primer matching the input sequence (16S-V4-R). Only dumbbells, and not unmodified inputs, yielded positive bands. SL and SL10Ta were also used for Phi29-based rolling circle amplification^2^ (**Fig. 2b**). Only dumbbells, and not unmodified inputs, allowed Phi29-catalyzed polymerization of high molecular-weight products (**Fig. 2b**, arrow).

### Tunneling to PacBio Sequencing

DNA dumbbells are the template format of PacBio sequencers. Analyses of a standard microbial community on the PacBio platform are published in Callahan et al^10^. For comparison, we prepared a full-length 16S library from the same standard community with customized PCR primers (**Fig. 3a and Supplementary Table 1**), along with other full-length 16S control libraries (**Supplementary Table 2**). These PCR oligos differed from PacBio standards on the 5’ ends, where the forward primer had Illumina P5 sequences and the reverse primer had Illumina P7 sequences, respectively. After PCR amplification, these linear DNA libraries were converted to dumbbell derivatives.

These libraries were sequenced on the PacBio Sequel platform with default parameters. Demultiplexed reads of the standard community were analyzed with the pipeline from Callahan et al^10^. 23 amplicon sequence variants (ASV) of full-length 16S were identified by DADA2^11^, 12 of which exactly matched reference sequences (**Table 1**). All eight bacterial genera, including *Bacillus*, *Enterococcus*, *Escherichia*, *Lactobacillus*, *Listeria*, *Pseudomonas*, *Salmonella*, and *Staphylococcus* were detected.

**Table 1.** Matches of ASV to ZymoBIOMICS reference

| Genus | No | Yes |
| --- | --- | --- |
| *Bacillus* | 0 | 3 |
| *Enterococcus* | 0 | 1 |
| *Escherichia* | 5 | 0 |
| *Lactobacillus* | 5 | 0 |
| *Listeria* | 0 | 3 |
| *Pseudomonas* | 0 | 1 |
| *Salmonella* | 0 | 2 |
| *Staphylococcus* | 1 | 2 |

Their genomic abundances are expected to be equal. Our results recapitulated the prediction with a relatively even distribution (**Fig. 3b**). Precise stoichiometry of ASV per genome (**Fig. 3c**) corresponding to Callahan’s results^10^ was successfully reproduced.

Twelve fecal full-length 16S libraries previously sequenced following PacBio’s standard protocol (**Supplementary Table 2**), henceforth referred to as direct sequencing (DS), were re-sequenced as controls. ASV of each sample by DS and tunneled sequencing (TS) were both identified by DADA2^11^. Although DS had higher HiFi read depths than TS (14.4 fold on average), ASV identified from DS were only 4.5 fold higher than TS (**Table 2**). Among TS ASV, 92.2±2.1% (s.e.) overlapped with those from DS. Abundances of overlapped ASV showed significant correlations (**Fig. 3d**). Pearson coefficients ranged from 0.77 to 0.99 (0.89±0.02, s.e.), with a significance level <10^-6^ at least.

**Table 2.**
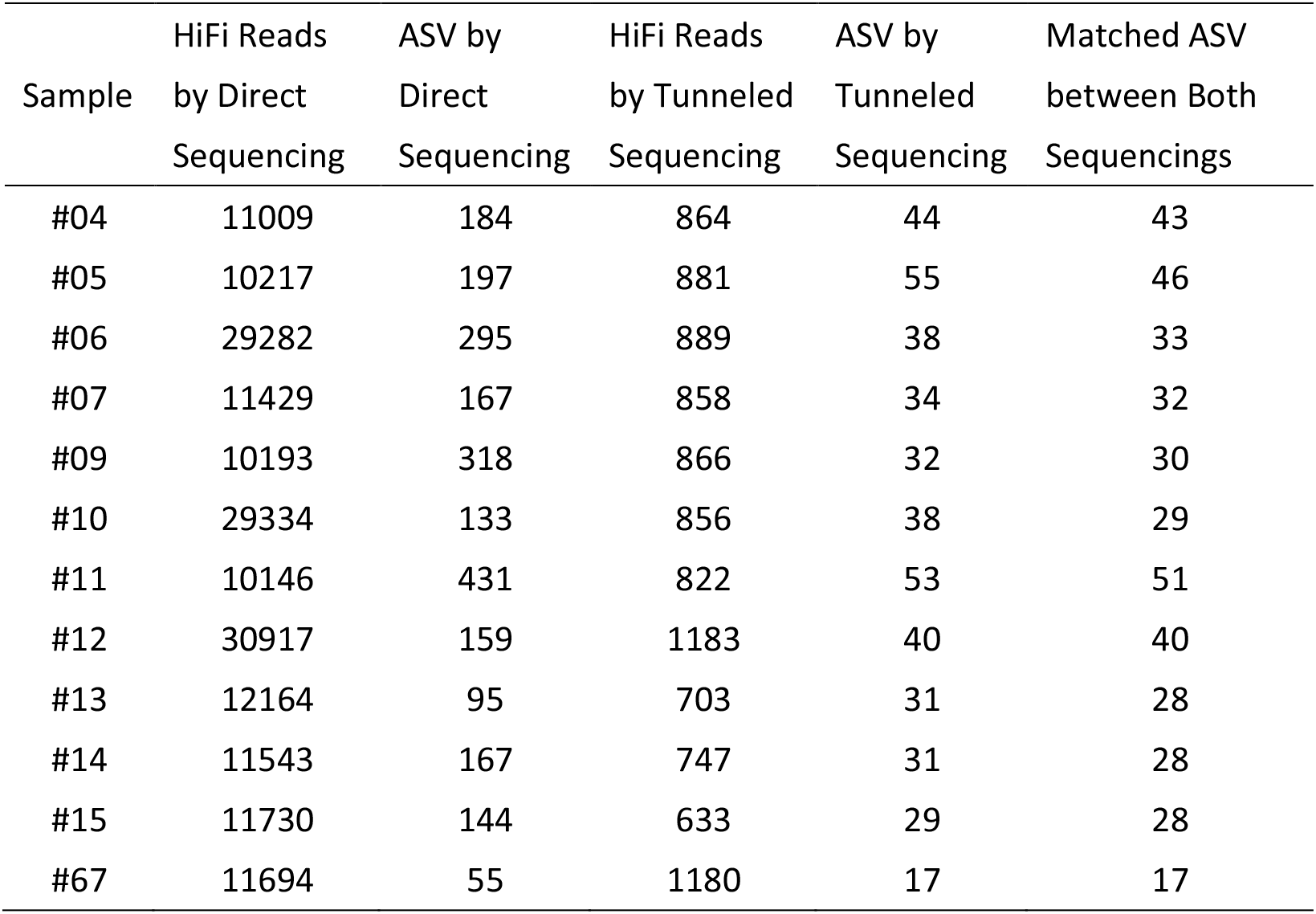
Summary of HiFi reads and ASV from direct and tunneled sequencing

| Sample | HiFi Reads<br>by Direct<br>Sequencing | ASV by<br>Direct<br>Sequencing | HiFi Reads<br>by Tunneled<br>Sequencing | ASV by<br>Tunneled<br>Sequencing | Matched ASV<br>between Both<br>Sequencings |
| --- | --- | --- | --- | --- | --- |
| #04 | 11009 | 184 | 864 | 44 | 43 |
| #05 | 10217 | 197 | 881 | 55 | 46 |
| #06 | 29282 | 295 | 889 | 38 | 33 |
| #07 | 11429 | 167 | 858 | 34 | 32 |
| #09 | 10193 | 318 | 866 | 32 | 30 |
| #10 | 29334 | 133 | 856 | 38 | 29 |
| #11 | 10146 | 431 | 822 | 53 | 51 |
| #12 | 30917 | 159 | 1183 | 40 | 40 |
| #13 | 12164 | 95 | 703 | 31 | 28 |
| #14 | 11543 | 167 | 747 | 31 | 28 |
| #15 | 11730 | 144 | 633 | 29 | 28 |
| #67 | 11694 | 55 | 1180 | 17 | 17 |

## Discussion

In this study, we demonstrate a simple method to convert P5/P7-ended short-read libraries into DNA dumbbells in one step. Resulting dumbbells are ready for Phi29-based rolling circle amplification (RCA)^2^. PacBio uses RCA as its *bona fide* sequencing mechanism. Oxford Nanopore can employ RCA to increase sequencing accuracy^12, 13^. MGI Tech uses RCA to prepare sequencing templates as ‘DNA nanoballs’^4^. In other words, our method provides a unified one-step option for tunneling any P5/P7-ended sequencing library to a long-read format suitable for many different sequencers.

Previous methods for forming DNA dumbbells use multiple steps and employ PCR^14, 15^. Our method instead combines the principle of Gibson Assembly^9^ and the unique topology of self-looping oligonucleotides to make DNA dumbbells in one step. This protocol modifies only the end nucleotides of input libraries. Native nucleotides inside flanking P5/P7 sequences can in principle be left untouched. Asymmetric dumbbells can be readily achieved by ligating P5- or P7-specific loops. Methods for linking multiple dumbbells in tandem can be pursued with minor modifications.

We demonstrated the tunneling option on a PacBio sequencer (**Fig. 3**). Although read depths were suboptimal (**Table 2**), analyses of the standard microbial community exhibited very good correspondence to previously published results^10^. We saw reasonably distributed abundances of bacterial genomes from the mock standard community. In total, 23 full-length 16S ASV were identified. The number was six fewer than identified by Callahan et al^10^. Due to the significant difference in sequencing depths, this disparity is expected. Importantly, stoichiometry of ASV per genome matched Callahan’s results exactly. The missing three ASV for *Bacillus,* two ASV for *Enterococcus,* and one ASV for *Staphylococcus* did not change overall integral sums of ASV for these genera, which remained 10, 4, and 6, respectively. Even for fecal samples with complex bacterial communities, reads of tunneled libraries correlated well (**Fig. 3d**).

The reason for suboptimal read depths from tunneled sequencing on the PacBio platform is unknown. Potential explanations include sequencing primers mismatching our customized loop, unoptimized stoichiometry of sequencing primers *vs.* asymmetric dumbbell templates, and inefficient docking of the primer-dumbbell complex. All of these can be corrected with fine-tuning of our technique.

## Supporting information

Supplementary Tables and Bioinformatic Workflow

## Acknowledgements

We thank Chih-Wei Joshua Liu (MIT Physics of Living Systems) for English editing. Human samples previously sequenced on PacBio sequencers were kindly provided by Dr. Yu-Chen Lin at Far-Eastern Memorial Hospital. This work was supported by National Health Research Institutes (MG-109-SP-05).

## Author Contributions

Y.H.C. conducted experiments. H.H.L. developed the method.

## Competing Financial Interests

A patent application is being filed.

## Notes

### Summary of Updates

'Pacific Biotechnology' corrected to 'Pacific Biosciences' in texts; Figure 3 updated to include new results; New tables added; Supplemental file updated

