## Supplementary Tables and Bioinformatic Workflow for "A One-Step Synthesis of DNA Dumbbells Tunnels Short-Read Libraries into a Long-Read Format"

**Supplementary Table 1** Primer sequences for tunneled PacBio sequencing

| Name | Sequences[1] |
|---|---|
| 16S-F-bc1005-P5 | 5'-AATGATACGGCGACCACCGAGATCTACAC **CACTCGACTCTCGCGT** AGRGTTYGATYMTGGCTCAG-3' |
| 16S-F-bc1007-P5 | 5'-AATGATACGGCGACCACCGAGATCTACAC **TCTGTATCTCTATGTG** AGRGTTYGATYMTGGCTCAG-3' |
| 16S-F-bc1020-P5 | 5'-AATGATACGGCGACCACCGAGATCTACAC **CACGACACGACGATGT** AGRGTTYGATYMTGGCTCAG-3' |
| 16S-R-bc1033-P7 | 5'-CAAGCAGAAGACGGCATACGAGAT **AGAGACTGCGACGAGA** RGYTACCTTGTTACGACTT-3' |
| 16S-R-bc1035-P7 | 5'-CAAGCAGAAGACGGCATACGAGAT **CAGAGAGTGCGCGCGC** RGYTACCTTGTTACGACTT-3' |
| 16S-R-bc1044-P7 | 5'-CAAGCAGAAGACGGCATACGAGAT **CGCGCGTCGTCTCAGC** RGYTACCTTGTTACGACTT-3' |
| 16S-R-bc1045-P7 | 5'-CAAGCAGAAGACGGCATACGAGAT **AGAGAGTACGATATGT** RGYTACCTTGTTACGACTT-3' |
| 16S-R-bc1054-P7 | 5'-CAAGCAGAAGACGGCATACGAGAT **TCTGTAGTGCGTGCGC** RGYTACCTTGTTACGACTT-3' |
| 16S-R-bc1056-P7 | 5'-CAAGCAGAAGACGGCATACGAGAT **ATGTGCGTGTGTGTCT** RGYTACCTTGTTACGACTT-3' |
| 16S-R-bc1057-P7 | 5'-CAAGCAGAAGACGGCATACGAGAT **CTCTCAGACGCTCGTC** RGYTACCTTGTTACGACTT-3' |
| 16S-R-bc1060-P7 | 5'-CAAGCAGAAGACGGCATACGAGAT **TGTGTCTATACTCATC** RGYTACCTTGTTACGACTT-3' |
| 16S-R-bc1062-P7 | 5'-CAAGCAGAAGACGGCATACGAGAT **TATAGACTATCTGAGA** RGYTACCTTGTTACGACTT-3' |
| 16S-R-bc1065-P7 | 5'-CAAGCAGAAGACGGCATACGAGAT **GTATGTGAGAGAGCGC** RGYTACCTTGTTACGACTT-3' |
| 16S-R-bc1075-P7 | 5'-CAAGCAGAAGACGGCATACGAGAT **CACGCGACGCTCTCTA** RGYTACCTTGTTACGACTT-3' |

[1] Barcodes are in **bold**.

**Supplementary Table 2** Summary of samples for tunneled PacBio sequencing

| Sample | HiFi Reads | Notes | Asymmetric Loops[1] |
|---|---|---|---|
| #01 | 1243 | DH5$\alpha$ | Yes |
| #02 | 1693 | DH5$\alpha$ | No |
| #03 | 1107 | ZymoBIOMICS standard | Yes |
| #04 | 864 | PacBio sequenced sample | Yes |
| #05 | 881 | PacBio sequenced sample | Yes |
| #06 | 889 | PacBio sequenced sample | Yes |
| #07 | 858 | PacBio sequenced sample | Yes |
| #09 | 866 | PacBio sequenced sample | Yes |
| #10 | 856 | PacBio sequenced sample | Yes |
| #11 | 822 | PacBio sequenced sample | Yes |
| #12 | 1183 | PacBio sequenced sample | Yes |
| #13 | 703 | PacBio sequenced sample | Yes |
| #14 | 747 | PacBio sequenced sample | Yes |
| #15 | 633 | PacBio sequenced sample | Yes |
| #67 | 1180 | PacBio sequenced sample | Yes |

[1] Asymmetric loops allow hybridization of the sequencing primer only on the loop at the P5 end but not on the loop at the P7 end.

**Bioinformatic workflow to analyze ZymoBIOMICS reference sample after tunneled sequencing**

**# At Bash shell**

```
$ cat adapters.fasta
>p5
CGTATCATCTCTCTCTTTTCCTCCTCCTCCGTTGTTGTTGTTGAGAGAGATGATACG
>p7
GAAGTGCGCTGTAAGTATTACCAGCGCACTTC
$ recalladapters –o readapt ––adapters ./adapters.fasta –s subreadset.xml
$ ccs readapt.subreads.bam readapt.fastq.gz
```

**# At ipython3**

```
import gzip
from Bio import SeqIO

with gzip.open("readapt.fastq.gz","rt") as handle:
       readapt = list(SeqIO.parse(handle, "fastq"))

import numpy as np

# size selection [1000, 2000]
fl = list(filter(lambda i: len(i)>=1000 and len(i)<=2000, readapt))

# demultiplex
p5 = 'ATGATACGGCGACCACCGAGATCTACAC'
p7 = 'CAAGCAGAAGACGGCATACGAGAT'

from Bio import Seq

n=19
p5Seed = p5[-n:]
p7Seed = p7[-n:]

from fuzzywuzzy import fuzz
```

```
from scipy.signal import find_peaks


r = 50  # length of adapter scan region

h = 90  # peak height threshold


# to find p5Seed peak at 5'end

seed = p5Seed

targets = map(lambda s: seed + str(s.seq[:r]), fl)  #find_peaks cannot identify peak at position 0, so pad a beginning seed

ratiosArray = map(lambda tandem: np.array(list(map(lambda i: fuzz.ratio(seed, tandem[i:i+n]), range(len(tandem)-n)))), targets)

p5BCIndex = list(map(lambda ratios: find_peaks(ratios, height=h)[0], ratiosArray))


# p5Seed rev_comp at 3'end

targets = map(lambda s: seed + str(s.seq[-r:].reverse_complement()), fl)

ratiosArray = map(lambda tandem: np.array(list(map(lambda i: fuzz.ratio(seed, tandem[i:i+n]), range(len(tandem)-n)))), targets)

p5RCBCIndex = list(map(lambda ratios: find_peaks(ratios, height=h)[0], ratiosArray))


# to find p7Seed peak at 5'end

seed = p7Seed

targets = map(lambda s: seed + str(s.seq[:r]), fl)  #find_peaks cannot identify peak at position 0, so pad a beginning seed

ratiosArray = map(lambda tandem: np.array(list(map(lambda i: fuzz.ratio(seed, tandem[i:i+n]), range(len(tandem)-n)))), targets)

p7BCIndex = list(map(lambda ratios: find_peaks(ratios, height=h)[0], ratiosArray))


# p7Seed rev_comp at 3'end

targets = map(lambda s: seed + str(s.seq[-r:].reverse_complement()), fl)

ratiosArray = map(lambda tandem: np.array(list(map(lambda i: fuzz.ratio(seed, tandem[i:i+n]), range(len(tandem)-n)))), targets)

p7RCBCIndex = list(map(lambda ratios: find_peaks(ratios, height=h)[0], ratiosArray))


bc = 16   # barcode length
```

```
def binning(index):
	p5f = p5BCIndex[index]
	p5r = p5RCBCIndex[index]
	p7f = p7BCIndex[index]
	p7r = p7RCBCIndex[index]
	fasta = str(fl[index].seq)
	anchorL = 0
	anchorR = 0
	rev = False
	if p5f.size:
		p5bc = fasta[p5f[0]:p5f[0]+bc]
		if p7r.size:
			p7bc = str(Seq.Seq(fasta[-p7r[0]-bc:-p7r[0]]).reverse_complement())
			anchorL = p5f[0]+bc
			anchorR = -p7r[0]-bc
		else:
			p7bc = ''
	else:
		p5bc = ''
		if p7f.size:
			p7bc = fasta[p7f[0]:p7f[0]+bc]
			if p5r.size:
				p5bc = str(Seq.Seq(fasta[-p5r[0]-bc:-p5r[0]]).reverse_complement())
				rev = True
				anchorL = p7f[0]+bc
				anchorR = -p5r[0]-bc
		else:
			p7bc = ''
	return (p5bc, p7bc, anchorL, anchorR, rev)


flBin = list(map(lambda i: binning(i), range(len(fl))))


# BCDic
```

```
BCDic = {
"CACTCGACTCTCGCGT":"bc1005",
"TCTGTATCTCTATGTG":"bc1007",
"CACGACACGACGATGT":"bc1020",
"AGAGACTGCGACGAGA":"bc1033",
"CAGAGAGTGCGCGCGC":"bc1035",
"CGCGCGTCGTCTCAGC":"bc1044",
"AGAGAGTACGATATGT":"bc1045",
"TCTGTAGTGCGTGCGC":"bc1054",
"ATGTGCGTGTGTGTCT":"bc1056",
"CTCTCAGACGCTCGTC":"bc1057",
"TGTGTCTATACTCATC":"bc1060",
"TATAGACTATCTGAGA":"bc1062",
"GTATGTGAGAGAGCGC":"bc1065",
"CACGCGACGCTCTCTA":"bc1075"
}

sampleDic = {
("bc1005","bc1033"):"#01",
("bc1005","bc1035"):"#02",
("bc1005","bc1044"):"#03",
("bc1005","bc1045"):"#04",
("bc1005","bc1054"):"#05",
("bc1005","bc1056"):"#06",
("bc1005","bc1057"):"#07",
("bc1005","bc1060"):"#09",
("bc1005","bc1062"):"#10",
("bc1005","bc1065"):"#11",
("bc1005","bc1075"):"#12",
("bc1007","bc1033"):"#13",
("bc1007","bc1035"):"#14",
("bc1007","bc1044"):"#15",
("bc1020","bc1057"):"#67"
}
```

```
def binSample(p5bc, p7bc):
    try:
        sample = sampleDic[(BCDic[p5bc], BCDic[p7bc])]
    except:
        sample = "unknown"

    return sample

demulti = list(map(lambda x: binSample(x[0], x[1]), flBin))

# To prepare ZYMO fastq
indices = filter(lambda i: demulti[i] == '#03', range(len(demulti)))
with gzip.open("zymo_CCS_99_9.fastq.gz","wt") as handle:
    list(map(lambda i: 
handle.write(">"+fl[i][flBin[i][2]:flBin[i][3]].format("fastq")), indices))
```

**# At R**

The analytic instructions follow https://benjjneb.github.io/LRASManuscript/LRASms_Zymo.html.
